# Ophthalmic Changes in a Spaceflight Analog Are Associated with Brain Functional Reorganization

**DOI:** 10.1101/2020.09.09.289827

**Authors:** Heather R. McGregor, Jessica K. Lee, Edwin R. Mulder, Yiri E. De Dios, Nichole E. Beltran, Igor S. Kofman, Jacob J. Bloomberg, Ajitkumar P. Mulavara, Scott M. Smith, Sara R. Zwart, Rachael D. Seidler

**Author notes:** Corresponding author: Rachael D. Seidler, Ph.D., University of Florida, 1864 Stadium Rd., Gainesville, FL 32611. **Author Contributions**: HRM analyzed fMRI data. JKL collected data. JKL, YED, NEB, ISK, RDS set up the experiment. SRZ and SMS analyzed genetic and biochemical data. RDS, ERM, JJB, APM designed the experiment and secured funding. HRM drafted the manuscript. HRM, JKL, ERM, JJB, APM, SMS, SRZ, RDS edited the manuscript. **Role of the Funder/Sponsor**: The funding sources had no role in the design and conduct of the study; collection, management, analysis, and interpretation of the data; preparation, review, or approval of the manuscript; and decision to submit the manuscript for publication. **Conflict of Interest Disclosures**: None. **Data Availability Statement**: MRI files for this study will be placed in the NASA data repository upon study completion.

## Abstract

Following long-duration spaceflight, some astronauts exhibit ophthalmic structural changes referred to as Spaceflight Associated Neuro-ocular Syndrome (SANS). Optic disc edema is a common sign of SANS. The origin and effects of SANS are not understood as signs of SANS have not manifested in previous spaceflight analog studies. In the current spaceflight analog study, eleven subjects underwent 30 days of strict head down-tilt bed rest in elevated ambient carbon dioxide (HDBR+CO_2_). Using functional magnetic resonance imaging (fMRI), we acquired resting-state fMRI data at 6 time points: before (2), during (2), and after (2) the HDBR+CO_2_ intervention. Five participants developed optic disc edema during the intervention (SANS subgroup) and 6 did not (NoSANS group). This occurrence allowed us to explore whether development of signs of SANS during the spaceflight analog impacted resting-state functional connectivity during HDBR+CO_2_. In light of previous work identifying genetic and biochemical predictors of SANS, we further assessed whether the SANS and NoSANS subgroups exhibited differential patterns of resting-state functional connectivity prior to the HDBR+CO_2_ intervention. We found that the SANS and NoSANS subgroups exhibited distinct patterns of resting-state functional connectivity changes during HDBR+CO_2_ within visual and vestibular-related brain networks. The SANS and NoSANS subgroups also exhibited different resting-state functional connectivity prior to HDBR+CO_2_ within a visual cortical network and within a large-scale network of brain areas involved in multisensory integration. We further present associations between functional connectivity within the identified networks and previously identified genetic and biochemical predictors of SANS. Subgroup differences in resting-state functional connectivity changes may reflect differential patterns of visual and vestibular reweighting as optic disc edema develops during the spaceflight analog. This finding suggests that SANS impacts not only neuro-ocular structures, but also functional brain organization. Future prospective investigations incorporating sensory assessments are required to determine the functional significance of the observed connectivity differences.

**HIGHLIGHTS:** We investigated resting-state functional connectivity (FC) during a spaceflight analog with elevated CO_2_ (HDBR+CO_2_).

During the HDBR+CO_2_ intervention, a subset of participants developed optic disc edema, a sign of spaceflight-associated neuro-ocular syndrome (SANS).

Participants with signs of SANS exhibited a distinct pattern of resting-state functional connectivity changes within visual and vestibular-related networks during HDBR+CO_2_.

Participants who developed optic disc edema exhibited different FC prior to the spaceflight analog within a visual cortical network and within a large-scale network of brain areas involved in multisensory integration.

## 1. INTRODUCTION

As NASA prepares for the first human mission to Mars, it is critical to understand how spaceflight affects the brain and body. During long-duration missions onboard the International Space Station, some astronauts develop ophthalmic abnormalities including optic disc edema, choroidal folds, cotton wool spots, posterior globe flattening, and distention of the optic nerve sheath ^1–3^. These findings have been collectively termed Spaceflight Associated Neuro-ocular Syndrome (SANS). The origin and effects of SANS are not understood, and present significant potential risk to astronaut health and performance. Here we explored whether the development of signs of SANS during a spaceflight analog is associated with altered brain functional organization.

The effects of spaceflight are often studied using ground-based analogs, with head down-tilt bed rest (HDBR) being one of the most widely used ^4,5^. The head down-tilt posture simulates microgravity effects on the body including headward fluid shifts and axial body unloading. Headward fluid shift is hypothesized to be a contributing factor of SANS ^1^. Spaceflight has been shown to alter cerebrospinal fluid hydrodynamics ^6^, and result in ventricular volume increases ^6–8^, which have been associated with ocular globe deformation ^7^ and visual acuity visual changes (i.e., hyperopic refractive error shifts) ^8^. However, until recently, signs of SANS have not manifested in any spaceflight analog ^9^. Another potential contributing factor to SANS development during spaceflight is the mild hypercapnic environment of the International Space Station ^10^ where CO_2_ levels (0.5%) are more than 10 times higher than on Earth ^11^. As CO_2_ is a potent arterial vasodilator, small increases in arterial CO_2_ increase cerebral blood flow and intracranial blood volume ^12–14^, which can elevate intracranial pressure. As such, microgravity-induced headward fluid shifts combined with exposure to elevated CO_2_ may contribute to SANS development during spaceflight.

To better replicate the International Space Station environment, NASA commissioned the current spaceflight analog study employing 30 days of strict HDBR in elevated ambient CO_2_ (HDBR+CO_2_). The ambient CO_2_ was elevated to 0.5% during HDBR to match the mild hypercapnic environment of the International Space Station ^11^. This study used “strict” HDBR such that subjects remained in the 6° head down-tilt position at all times, and were not permitted to use a standard pillow ^15^. To investigate the brain’s adaptation to the HDBR+CO_2_ spaceflight analog, we assessed resting-state brain activity before, during, and after bed rest using functional magnetic resonance imaging (fMRI). Using this same dataset, we recently reported resting-state connectivity changes throughout the HDBR+CO_2_ intervention across the entire group as well as brain-behavior associations^16^. We reported that exposure to the HDBR+CO_2_ intervention resulted in gradual resting-state functional connectivity (FC) changes involving vestibular, visual, somatosensory and motor brain areas. During the HDBR+CO_2_ intervention, a subset of participants developed optic disc edema — a sign of SANS ^3,17,18^. This occurrence afforded us the unique opportunity to explore functional brain differences between participants who developed optic disc edema during the spaceflight analog (SANS subgroup) and those who did not (NoSANS subgroup). Here, we examined whether the SANS and NoSANS subgroups exhibited different longitudinal changes in FC from before, during, to after the HDBR+CO_2_ intervention.

Genetic and biochemical risk factors for optic disc edema development have recently been identified in the SANS subgroup in the current study ^18,19^ as well as in astronauts who develop SANS ^20,21^. Genetic risk factors include G alleles for the 5-methyltetrahydrofolate-homocysteine methyltransferase reductase (*MTRR*) A66G gene polymorphism and C alleles for the serine hydroxymethyltransferase 1 (*SHMT1*) 1420 gene polymorphism. Low levels of B vitamins (B6, B12, and folate) have also been found to predict optic disc edema development with the current HDBR+CO_2_ intervention ^18^ and with spaceflight ^20,21^. These findings may help explain the underlying risk for individuals who develop SANS compared to those who do not. In light of this, we further investigated FC differences between the SANS and NoSANS subgroups prior to bed rest. We present qualitative associations between identified FC differences between the SANS and NoSANS subgroups with participants’ B vitamin concentration and risk alleles.

Previous studies have examined changes in FC with exposure to HDBR in standard ambient air. These studies have demonstrated FC changes among brain areas involved in somatosensory, vestibular, and motor function as well as brain areas involved in multisensory integration, and regions within the default mode network ^5,22–26^. Based on the findings of these previous HDBR studies in ambient air ^5,22–26^, we hypothesized that brain areas related to multisensory integration, and regions within somatosensory, motor, vestibular, visual, and default mode networks would exhibit differential FC between the SANS and NoSANS subgroups. As this was the first HDBR study to induce signs of SANS, we also used an exploratory hypothesis-free approach.

Our analysis of longitudinal FC changes showed that the SANS subgroup exhibited a distinct pattern of resting-state FC changes involving visual and vestibular brain regions during the HDBR+CO_2_ intervention compared to the NoSANS subgroup. Further, the SANS and NoSANS subgroups exhibited differential resting-state brain activity prior to HDBR+CO_2_ within a visual cortical network and within a large-scale network of brain areas involved in multisensory integration. While preliminary, these findings suggest that the development of optic disc edema during this spaceflight analog is associated with functional brain reorganization.

## 2. METHODS

### 2.1 Participants

Twelve healthy adults enrolled in this study (6 males, 6 females). Participants were nonsmokers between 24 and 55 years of age with a body mass index ranging from 19 to 30 kg/m^2^. Exclusion criteria included any chronic illness and pain, elevated risk of thrombosis, chronic hypertension, diabetes, obesity, arthritis, hyperlipidemia, hepatitis, or metabolic bone diseases. Participants were also screened for any eye conditions that significantly impair vision. All participants passed psychological screening and an Air Force Class III equivalent physical examination. One female participant withdrew from the study on the first day of HDBR+CO_2_. Eleven participants in total completed the study; their demographics are summarized in Table 1. All participants provided written informed consent to study protocols approved by the ethics committee of the local medical association (Ärztekammer Nordrhein) and the institutional review boards at The University of Florida and NASA Johnson Space Center.

**Table1:** Participant demographics for the entire group (top) and for the SANS and NoSANS subgroups (bottom). Age is in years. SANS, spaceflight associated neuro-ocular syndrome; F, female; M, male.

|  | Group (n=11) |  |
| --- | --- | --- |
| Sex | 5 females | 6 males |
| Mean age (SD) | 33.9 (8.0) |  |
|  | SANS subgroup (n=5) | NoSANS subgroup (n=6) |
| Sex | 3 F, 2M | 2 F, 4 M |
| Mean age (SD) | 37.4 (7.6) | 30.3 (7.3) |

During the HDBR+CO_2_ intervention, 5 participants developed optic disc edema (“SANS subgroup”), and 6 participants did not (“NoSANS subgroup”). Development of optic disc edema did not result in measurable visual changes in the SANS subgroup. As reported previously ^27^, there was no statistically reliable age difference between participants in the SANS and NoSANS subgroups.

### 2.2 Study Design

This study was carried out at :envihab, the German Aerospace Center’s medical research facility in Cologne, Germany. This experiment was conducted as part of the VaPER (Vision Impairment and Intracranial Pressure and Psychological :envihab Research) bed rest campaign. Participants resided at the :envihab facility for the duration of this 58-day study. Figure 1 shows the study design and each phase is summarized below.

**Figure 1.**
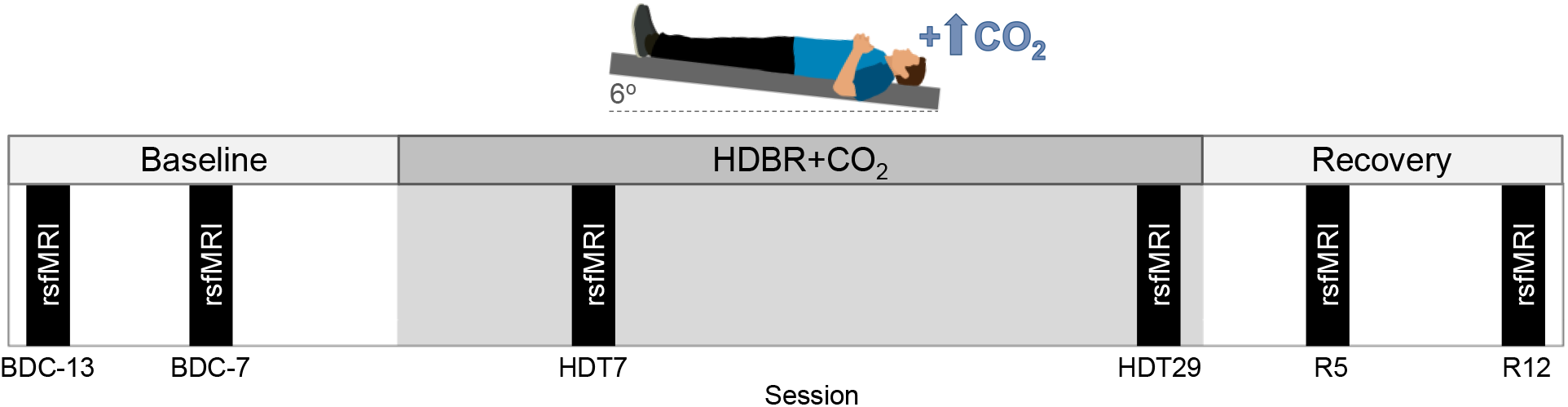
Baseline data collection (BDC) was carried out during a 14-day ambulatory phase in standard ambient air. Participants then underwent 30 days of strict 6° head down-tilt bed rest with 0.5% ambient carbon dioxide (HDBR+CO_2_). The bed rest intervention was followed by a 14-day ambulatory recovery (R) phase in standard ambient air. Resting-state fMRI scans were acquired on sessions BDC-13, BDC-7, HDT7, HDT29, R5, R12.

#### 2.2.1 Baseline data collection (BDC)

Baseline data collection (BDC) occurred during a 14-day phase (BDC-14 to BDC-1). Participants were ambulatory and the ambient air was standard (i.e., not hypercapnic). Participants remained inside the bed rest module of the :envihab facility, and their activity level was restricted to walking. Participants slept in a horizontal (i.e., 0° tilt) position. Resting-state fMRI scans were acquired twice during the baseline phase: 13 and 7 days before the bed rest phase (BDC-13 and BDC-7, respectively).

#### 2.2.2 Head down-tilt bed rest with elevated carbon dioxide (HDBR+CO_2_)

Following the BDC phase, participants underwent 30 days of strict 6° HDBR in elevated CO_2_ (HDBR+CO_2_). The ambient CO_2_ level was maintained at 0.5% during HDBR, corresponding to the average CO_2_ concentration onboard the ISS (3.8 mmHG ambient partial pressure of carbon dioxide)^11^. Participants adhered to bed rest 24 hours per day and were continuously monitored to ensure compliance. Participants were instructed to always have at least one of their shoulders in contact with the mattress. They were not permitted to raise or prop up their head at any time. We employed strict HDBR to keep the head at a −6° angle at all times and maintain elevated fluid pressure in the head ^15^. Participants were not permitted to use a standard pillow as previous research suggests that pillow use might confound results pertaining to SANS ^15^. Participants were provided a 5-cm tall neck support to use while lying on their side ^17,28^. Participants were restricted from raising or contracting their legs during HDBR+CO_2_. Physiotherapy and leg stretching was performed to avoid or alleviate muscle stiffness and back pain. Resting-state fMRI scans were acquired twice during bed rest: on days 7 and 29 of HDBR+CO_2_ (HDT7 and HDT29, respectively).

#### 2.2.3 Recovery (R)

Upon the completion of the HDBR+CO_2_ intervention, participants underwent a 14-day recovery phase (R+0 to R+13). As in the BDC phase, participants were ambulatory and the ambient air was standard (i.e., not hypercapnic). Participants remained inside the bed rest module of the :envihab facility, and their activity level was restricted to walking. Participants slept in a horizontal (i.e., 0° tilt) position. One-hour rehabilitation sessions took place 3, 5, 7, 8, 9, and 11 days following bed rest during which participants performed active stretching and exercises to improve strength and coordination using a coordination ladder or Bosu^®^ ball. Resting-state fMRI scans were acquired twice during the recovery phase: 5 and 12 days after the end of bed rest (R5 and R12, respectively).

### 2.3 MRI acquisition

For all six scan sessions, the participant’s body was in the 6° HDT position, but the head was horizontal in the MRI coil. A mask and tank system were used to maintain CO_2_ levels at 0.5% throughout MRI scans during the HDBR+CO_2_ phase. Neuroimaging data were acquired on the same 3T Siemens Biograph mMR scanner. The time of day of magnetic resonance imaging (MRI) scan sessions was held constant throughout the study, with all scans occurring in the morning.

T1-weighted anatomical images were acquired using a MPRAGE sequence (TR = 1900 ms, TE = 2.44 ms, flip angle = 9°, FOV = 250 × 250 mm, matrix = 512 × 512, voxel size = 0.5 × 0.5 mm, 192 sagittal slices, slice thickness = 1 mm, slice gap = 0.5 mm).

Whole-brain resting-state fMRI data were acquired using a T2*-weighted EPI sequence (TR = 2500 ms, TE = 32 ms, flip angle = 90°, FOV = 192 × 192 mm, matrix = 64 × 64, voxel size = 3 × 3 mm, 37 axial slices, slice thickness = 3.5mm, duration = 7 minutes). During resting-state scans, participants were instructed to remain awake and to fixate a red on-screen circle without thinking about anything in particular.

### 2.4 Image preprocessing

Image preprocessing has been detailed elsewhere ^16^. Neuroimaging data analyses were performed using the Statistical Parametric Mapping 12 software (SPM12; www.fil.ion.ucl.ac.uk/spm) and the CONN toolbox version 18a ^29^ implemented in Matlab 2018b. fMRI preprocessing followed a standard resting-state pipeline including slice timing correction, realignment, and coregistration of functional and anatomical images. Structural and functional images were segmented into gray matter, white matter, and cerebrospinal fluid (CSF) tissue maps, and then normalized to a Montreal Neurological Institute (MNI) standard space template.

To improve cerebellar normalization to MNI space, we extracted the cerebellum from each functional and anatomical image then normalized the cerebellums to the SUIT cerebellar template. To accomplish this, we used CERES to segment the cerebellum from each subject’s native space anatomical scan ^30^. CERES uses an automatic pipeline to perform cerebellar segmentation, and has been shown to yield more accurate segmentations than other toolboxes including SUIT ^31^. CERES-generated cerebellar masks were applied to coregistered T1 and functional images, allowing us to extract the cerebellum. The extracted cerebellums were then normalized to the SUIT cerebellar template. We chose to use the SUIT template because it shows greater anatomical detail within the cerebellum relative to whole brain MNI templates, yielding better normalization for fMRI data ^32^.

Whole-brain MRI-normalized functional images were smoothed using a 5 mm FWHM Gaussian kernel. MNI-normalized cerebellums extracted from functional scans were smoothed using a 2 mm FWHM Gaussian kernel. A smaller smoothing kernel was used to reduce smoothing across tissue types within the cerebellum’s smaller internal structures.

### 2.5 Image denoising

Head movement is known to influence measures of resting-state FC, introducing spurious correlations between distinct structures throughout the brain ^33,34^. We used the ARTifact detection tool (ART; http://www.nitrc.org/projects/artifact_detect), which is integrated into CONN, to detect and reject motion artifacts. Using CONN’s intermediate default motion threshold, a volume was considered an outlier if its composite movement exceeded 0.9 mm or if the global mean intensity of the volume exceeded 5 SD from the mean run image intensity. No more than 9 of the total 164 volumes (< 6%) were identified as outliers in any resting-state run. A split plot ANOVA followed by Bonferroni post-hoc correction revealed no significant subgroup x session interaction on the number of outlier volumes [Wilks’ F(2.46,22.11)=1.52, p=0.897], Greenhouse-Geisser corrected]. Similarly, there was no significant main effect of session on outlier volumes following Bonferroni post-hoc correction [Wilks’ F(5,5)=1.264, p=0.402].

Respiratory, pulsatile or cardiovascular noise also have confounding influences on resting-state fMRI data ^35,36^. To estimate and remove non-neuronal physiological noise, we used anatomical component-based noise correction (aCompCor) ^37^, which is integrated into CONN. For each session, the participant’s T1-weighted anatomical image was segmented into white matter and CSF masks, and then eroded by one voxel to reduce partial voluming effects. These masks were then applied to the unsmoothed resting-state data. Masks were applied to unsmoothed functional data so as to avoid signal being smoothed across different tissue types. The signal time courses were extracted from the white matter and CSF, and were used as noise regions of interest (ROIs) in a principal components analysis. Significant principal components were used to estimate the influence of noise as a voxel-wise linear combination of multiple estimated sources. The first five principal components were extracted for each of the white matter and CSF noise ROIs, and used as nuisance regressors in our first-level general linear model.

Preprocessed resting-state runs were denoised by regressing out confounding effects of head motion (6 head motion parameters and their 6 first-order temporal derivatives), 5 principal components modelling estimated noise from the white matter, 5 principal components modelling estimated noise from the CSF, and a “scrubbing” regressor modelling ART-detected outlier volumes within the run. Resting-state fMRI images were then band-pass filtered between 0.008–0.09 Hz ^38,39^.

### 2.6 Subject-level analyses

#### 2.6.1 Hypothesis-driven seed-to-voxel analyses

We used a hypothesis-driven approach to examine resting-state functional connectivity (FC) between 14 regions of interest (ROIs) and the rest of the brain. Drawing upon the findings of previous HDBR studies in standard ambient air, we hypothesized that the two subgroups would exhibit differential patterns of FC involving brain areas related to multisensory integration, somatosensation, motor function, vestibular processing, and regions within the default mode network ^5,24,25,40–43^. The coordinates for the ROIs were the same as those used in our previous resting-state fMRI study using HDBR intervention in standard ambient air ^5^. We also included ROIs located within left frontal eye field and bilateral primary visual cortices, the coordinates of which were determined using the automatic anatomic labeling (AAL) atlas included in the CONN toolbox. ROIs were 8-mm diameter spheres centered on coordinates presented in Table 2.

**Table 2.** Region of interest (ROI) coordinates used for hypothesis-based seed-to-voxel functional connectivity analyses.

| Regions of Interest (ROIs) | MNI Coordinates |  |  |
| --- | --- | --- | --- |
|  | *x* | *y* | *z* |
| Right premotor cortex (PMC) | 26 | -10 | 55 |
| Right middle frontal gyrus (MFG) | 30 | -1 | 43 |
| Left primary motor cortex (M1) | -38 | -26 | 50 |
| Left putamen | -28 | 2 | 4 |
| Left insular cortex | -41 | 12 | -9 |
| Right vestibular cortex | 42 | -24 | 18 |
| Left frontal eye field (FEF) | -27 | -9 | 64 |
| Left posterior cingulate cortex (PCC) | -12 | -47 | 32 |
| Right posterior parietal cortex (PPC) | 28 | -72 | 40 |
| Right superior posterior fissure | 24 | -81 | -36 |
| Left primary visual cortex (V1) | -10 | -70 | 10 |
| Right primary visual cortex (V1) | 12 | -66 | 10 |
| Right cerebellar lobule V | 16 | -52 | -22 |
| Right cerebellar lobule VIII | 20 | -57 | -53 |

We computed the average time series of unsmoothed functional data within each ROI. Time series for ROIs located in the cerebrum were extracted from the whole brain unsmoothed functional images (source image). Time series for ROIs located in the cerebellum were extracted from the unsmoothed cerebellums extracted from functional images (source image). FC was estimated by computing the Pearson’s correlation coefficient between each ROI time series and that of every other voxel in the cerebrum (i.e., smoothed whole brain functional images) or within the cerebellum (i.e., smoothed cerebellums extracted from functional scans). Fisher Z-transformation was then used to improve the normality of the resulting correlation coefficients. This procedure was repeated for each participant’s six resting-state runs, yielding per-session Z score maps reflecting FC between each ROI and the rest of the brain.

#### 2.6.2 Hypothesis-free voxel-to-voxel analyses

Since signs of SANS have never manifested in previous spaceflight analog studies, we also employed a hypothesis-free approach to explore connectivity differences between the SANS and NoSANS subgroups. This voxel-to-voxel analysis does not require *a priori* ROI selection and as such allowed us to identify subgroup differences that may otherwise be missed by our seed-to-voxel approach using hypotheses based on the findings of HDBR studies in standard ambient air. For this approach, we estimated the intrinsic connectivity contrast, a measure reflecting the strength of connectivity between a given voxel and every other voxel in the brain ^44^. First, we computed Pearson’s correlation coefficients between the time course of each voxel with that of every other voxel in the brain (or within the cerebellum). We calculated the root mean square of each correlation coefficient and then performed a Fisher Z-transformation. This procedure was repeated for each participant’s six resting-state runs, yielding per-session Z score maps reflecting the strength (i.e., magnitude) of connectivity between each voxel and the rest of the brain.

### 2.7 Group-level analyses

#### 2.7.1 Subgroup Differences in Longitudinal Changes in Functional Connectivity

We first examined if the SANS and NoSANS subgroups exhibited differential patterns of FC changes from before, during, to after the HDBR+CO_2_ intervention. We performed this longitudinal analysis using both hypothesis-based and hypothesis-free approaches. In our general linear model (GLM), subgroup was used as a between-subjects factor (i.e., SANS, NoSANS) and session was used as a within-subjects factor (i.e., 6 time points). The longitudinal within-subject contrast model used for this analysis was informed by the ophthalmological findings of another research group that participated in this HDBR+CO_2_ study ^3^. Laurie and colleagues measured total retinal thickness in these same participants using optical coherence tomography on days 1, 15, and 30 of the HDBR+CO_2_ intervention, and on the 6th and 13th day of the recovery phase. The SANS subgroup showed a gradual increase in total retinal thickness during HDBR+CO_2_ which was followed by a gradual post-bed rest decrease towards baseline values ^3^. Our longitudinal analysis aimed to identify brain networks in which FC changes mirrored the time course of optic disc edema onset and recovery. We therefore used a within-subject contrast model (shown in Figure 2) to identify brain areas in which FC was stable across the baseline phase (BDC-13, BDC-7) and then showed a linear FC change throughout HDBR+CO_2_ followed by a gradual return towards baseline in the subsequent recovery phase. Participant age and sex were included as covariates. The significance level was set at an uncorrected voxel threshold of *p* < 0.001 with a cluster-size threshold of *p* < 0.05 (two-tailed) corrected for multiple comparisons according to the false discovery rate (FDR) method.

**Figure 2.**
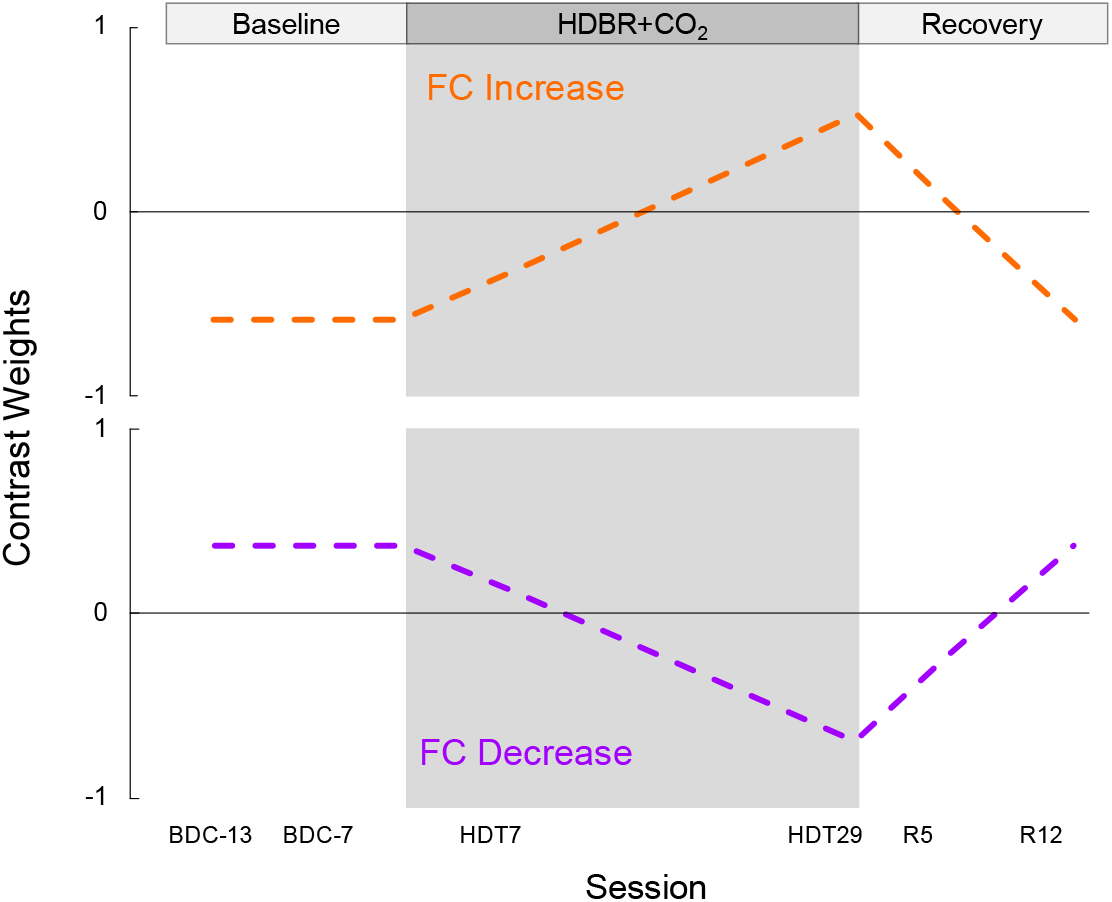
Contrast models used to longitudinal FC changes. The gray shaded region indicates the HDBR+CO_2_ phase. The model shown in orange was used to identify brain networks showing stable FC across the baseline phase followed by a gradual FC increase during HDBR+CO_2_, then a gradual return to baseline following bed rest. The model shown in purple was used to identify brain networks showing stable FC across the baseline phase followed by a gradual FC decrease during HDBR+CO_2_, then a gradual return to baseline following bed rest. BDC, baseline data collection; HDBR+CO_2_, head down-tilt bed rest with elevated carbon dioxide; R, recovery; FC, functional connectivity.

#### 2.7.2 Subgroup Differences in Functional Connectivity at Baseline

In light of recent work identifying genetic and biochemical predictors of optic edema development in these same participants ^3^, we also tested if there were FC differences between the SANS and NoSANS subgroups before bed rest. This analysis aimed to uncover functional brain networks in which resting-state activity was predictive of which participants would develop signs of SANS in the subsequent HDBR+CO_2_ intervention. For this analysis, our GLM included subgroup (i.e., SANS, NoSANS) as a between-subjects factor and baseline session (i.e., BDC-13, BDC-7) as a within-subjects factor. The within-subjects contrast modeled the average FC across the 2 BDC sessions. Participant age and sex were included as covariates. The significance level was set at an uncorrected voxel threshold of *p* < 0.001 with a cluster-size threshold of *p* < 0.05 (two-tailed) corrected for multiple comparisons according to the FDR method.

For each identified network by our baseline FC analysis, we performed a follow-up analysis to assess the test–retest reliability of FC values from BDC-13 to BDC-7. For each ROI-cluster connection, we computed the intraclass correlation coefficient between FC values on BDC-13 and on BDC-7 across all participants. This analysis, implemented using SPSS version 21 (SPSS Inc.), used a two-way mixed effects model to examine absolute agreement between the two FC measurements acquired during the baseline phase. Intraclass correlation coefficient values were classified into one of the following intervals: 0-0.2 (slight); 0.2-0.4 (fair); 0.4-0.6 (moderate); 0.6-0.8 (substantial); 0.8-1.0 (almost perfect) ^45^, with larger values indicating greater reliability of FC values between BDC sessions across participants.

A total of 30 tests were performed: 14 seed-to-voxel and 1 voxel-to-voxel analyses each for the longitudinal and baseline FC analyses. To control the false discovery rate, all cluster-size uncorrected p values were submitted to Benjamini and Hochberg’s FDR procedure using a *α*FDR of 0.05 ^46,47^. As a result, all clusters with uncorrected cluster size p < 0.0016 (i.e., (1/30) x 0.05) were deemed statistically significant.

### 2.8 Associations between Functional Connectivity and Genetic and B vitamin status

Biochemical and genetic data were collected from participants as part of NASA’s international standard measures assessments and other studies. A fasting blood sample was collected on BDC-11, 11 days before the HDBR+CO_2_ intervention. Samples were analyzed for *MTRR* 66 G alleles, *SHMT1* 1420 C alleles, B6 and B12 vitamin levels, and homocysteine as previously described ^18^. We quantified genetic risk for SANS by summing the number of *MTRR* 66 G alleles and *SHMT1* 1420 C alleles, as has been done in other studies ^18^.

For functional brain networks identified by our longitudinal analysis, we qualitatively examined associations between each risk factor and changes in FC from pre- to post-HDBR+CO_2_ (i.e., BDC-7 to HDT29). For functional brain networks identified by our baseline analysis, we qualitatively examined associations between each risk factor and average baseline FC values (i.e., across BDC-13 and BDC-7). As these genetic and biochemistry measures predict optic disc edema, which we used as a regressor (SANS vs. NoSANS) in our resting-state fMRI, it follows that FC differences in our identified networks should be related to risk factors. Due to the nonindependence of this analysis, we present associations for illustrative purposes only, not as the basis for inference ^48^.

## 3. RESULTS

### 3.1 Subgroup Differences in Longitudinal Changes in Functional Connectivity

Our hypothesis-based seed-to-voxel analyses revealed that the ROI in right posterior parietal cortex (PPC) and clusters within bilateral insular cortices exhibited differential longitudinal changes between the SANS and NoSANS subgroups (Figure 3A). The NoSANS subgroup showed decreased FC between these regions during HDBR+CO_2_ followed by a post-bed rest FC increase. In contrast, the SANS subgroup showed the opposite pattern, increasing FC during HDBR+CO_2_ followed by a post-bed rest decrease. Cluster coordinates and statistics are presented in Table 3.

**Figure 3.**
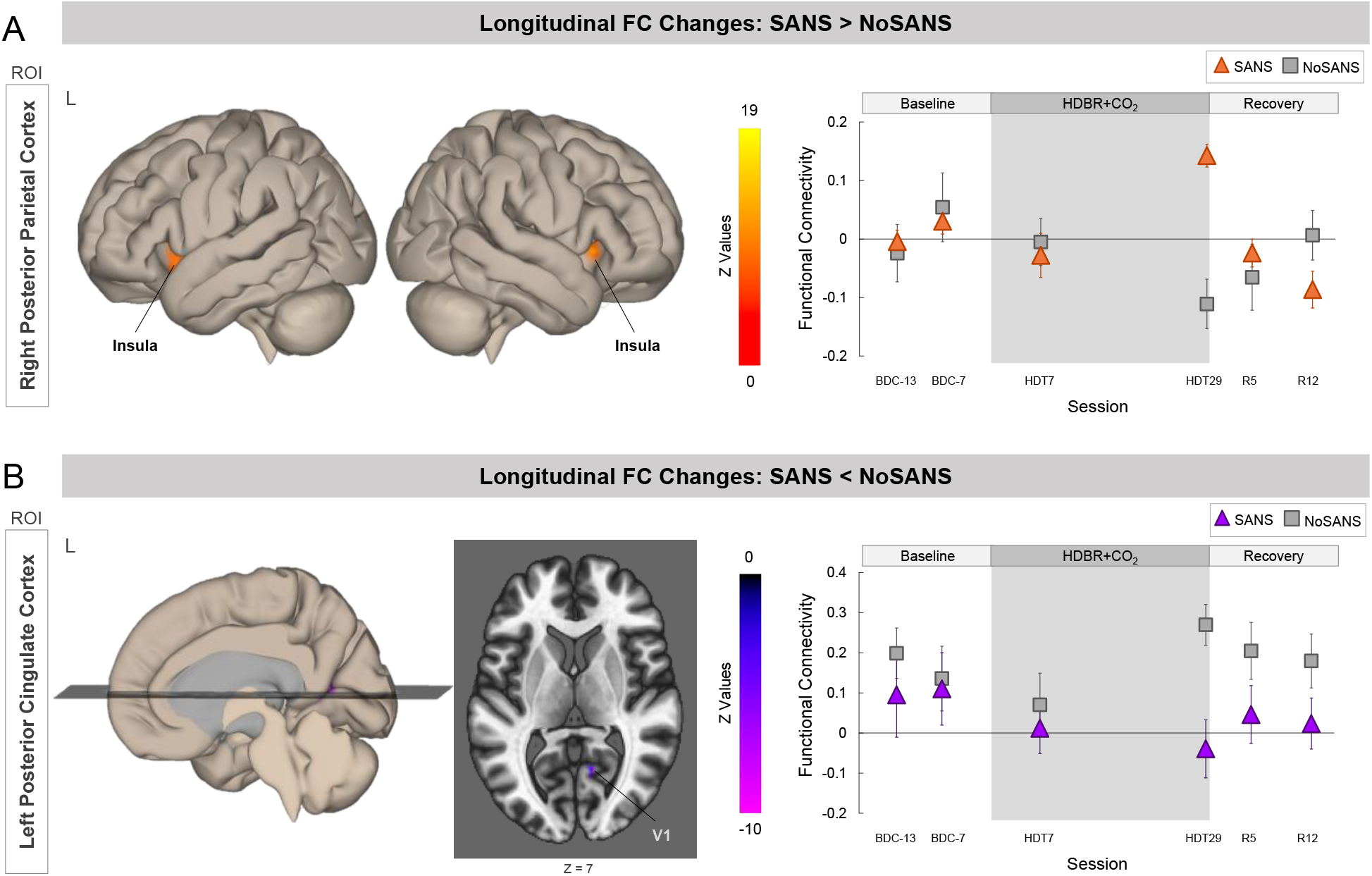
Differential longitudinal FC changes between the SANS and NoSANS subgroups. **A)** The NoSANS subgroup (gray squares) showed a FC decrease between the ROI in right posterior parietal cortex (PPC) and clusters in bilateral insular cortices during HDBR+CO_2_ followed by a post-bed rest reversal. The SANS subgroup (triangles) exhibited FC increases between these regions during HDBR+CO_2_ followed by a post-bed rest decrease. **B)** The NoSANS subgroup (gray squares) showed a FC increase between the ROI in left posterior cingulate cortex (PCC) and right primary visual cortex (V1) during HDBR+CO_2_ followed by a post-bed rest reversal. The SANS subgroup (triangles) exhibited a FC decrease between these regions during HDBR+CO_2_ followed by a post-bed rest increase. The dashed line on each graph represents the hypothesized longitudinal model. Error bars represent standard error. L, left; R, right; ROI, region of interest; FC, functional connectivity; SANS, spaceflight associated neuro-ocular syndrome; HDBR+CO_2_, head down-tilt bed rest with elevated CO_2_.

**Table 3:**
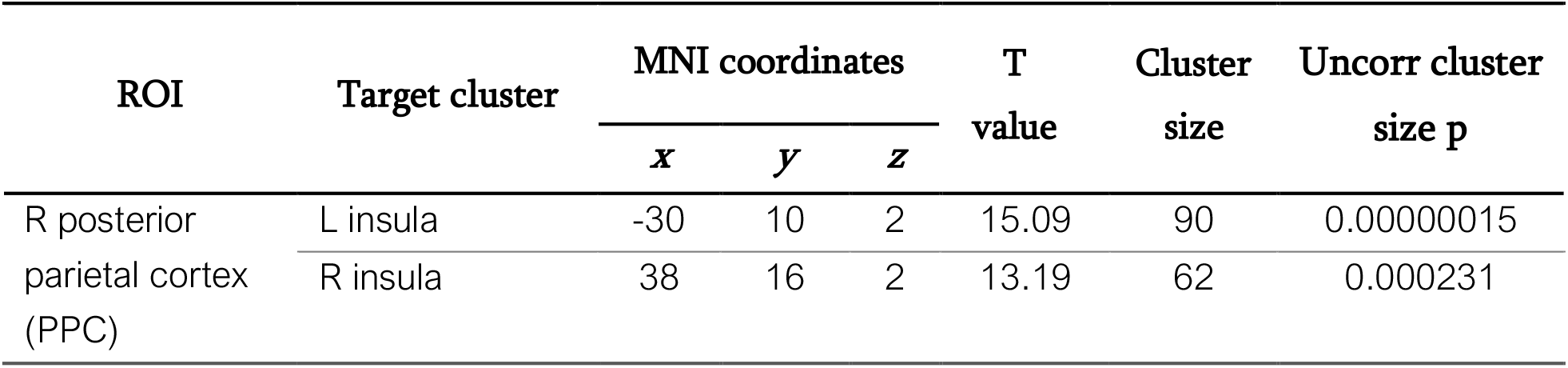
Longitudinal FC change results. Peak MNI coordinates and T value of the clusters in insular cortices. For the SANS subgroup, FC between the right PPC and bilateral insular cortices increased during HDBR+CO_2_ then decreased during the recovery phase. The NoSANS subgroup exhibited the opposite pattern. Cluster size indicates the number of voxels within each cluster. Uncorr cluster size p indicates uncorrected cluster-size p values. R, right; ROI, region of interest; FC, functional connectivity.

Furthermore, the ROI in left PCC exhibited differential longitudinal changes with right V1 between the two subgroups (Figure 3B). The NoSANS subgroup showed a FC increase between these regions during HDBR+CO_2_ which decreased after bed rest whereas the SANS subgroup showed decreased FC between these regions during HDBR+CO_2_ followed by a post-bed rest reversal. Cluster coordinates and statistics are presented in Table 4.

**Table 4:**
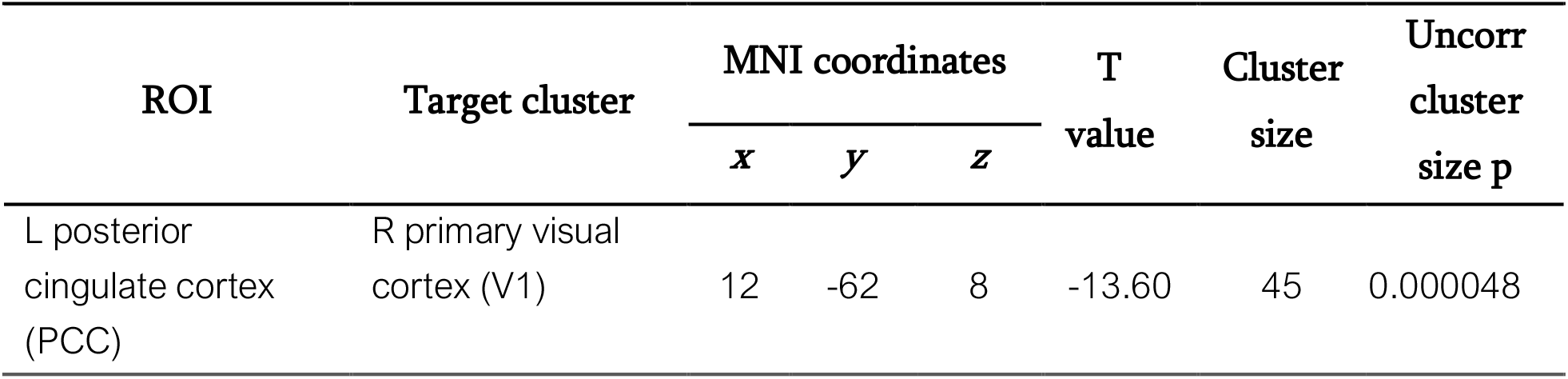
Longitudinal FC change results. Peak MNI coordinates and T value of the cluster within right V1. For the SANS subgroup, FC between PCC and V1 gradually decreased during HDBR+CO_2_ followed by a FC increase during the recovery phase. The NoSANS subgroup exhibited the opposite pattern. Cluster size indicates the number of voxels within each cluster. Uncorr cluster size p indicates uncorrected cluster-size p values. L, left; R, right; ROI, region of interest; FC, functional connectivity.

| ROI | Target cluster | MNI coordinates |  |  | T<br>value | Cluster<br>size | Uncorr<br>cluster<br>size p |
| --- | --- | --- | --- | --- | --- | --- | --- |
|  |  | *x* | *y* | *z* |  |  |  |
| L posterior<br>cingulate cortex<br>(PCC) | R primary visual<br>cortex (V1) | 12 | -62 | 8 | -13.60 | 45 | 0.000048 |

Our hypothesis-free voxel-to-voxel analysis did not yield any statistically significant clusters.

### 3.2 Subgroup Differences in Functional Connectivity at Baseline

We also identified two brain networks in which average FC across the baseline scan sessions significantly differed between the SANS and NoSANS subgroups. The first network, shown in Figure 4A, consisted of right primary visual cortex (V1) and higher order visual areas V3 and V4 within the left hemisphere. The NoSANS subgroup exhibited positive connectivity between these visual brain areas at baseline. In contrast, the SANS subgroup exhibited lower FC or an anti-correlation (i.e., negative FC) between right V1 and left V3 and V4 relative to the NoSANS subgroup. Cluster coordinates and statistics are presented in Table 5.

**Figure 4.**
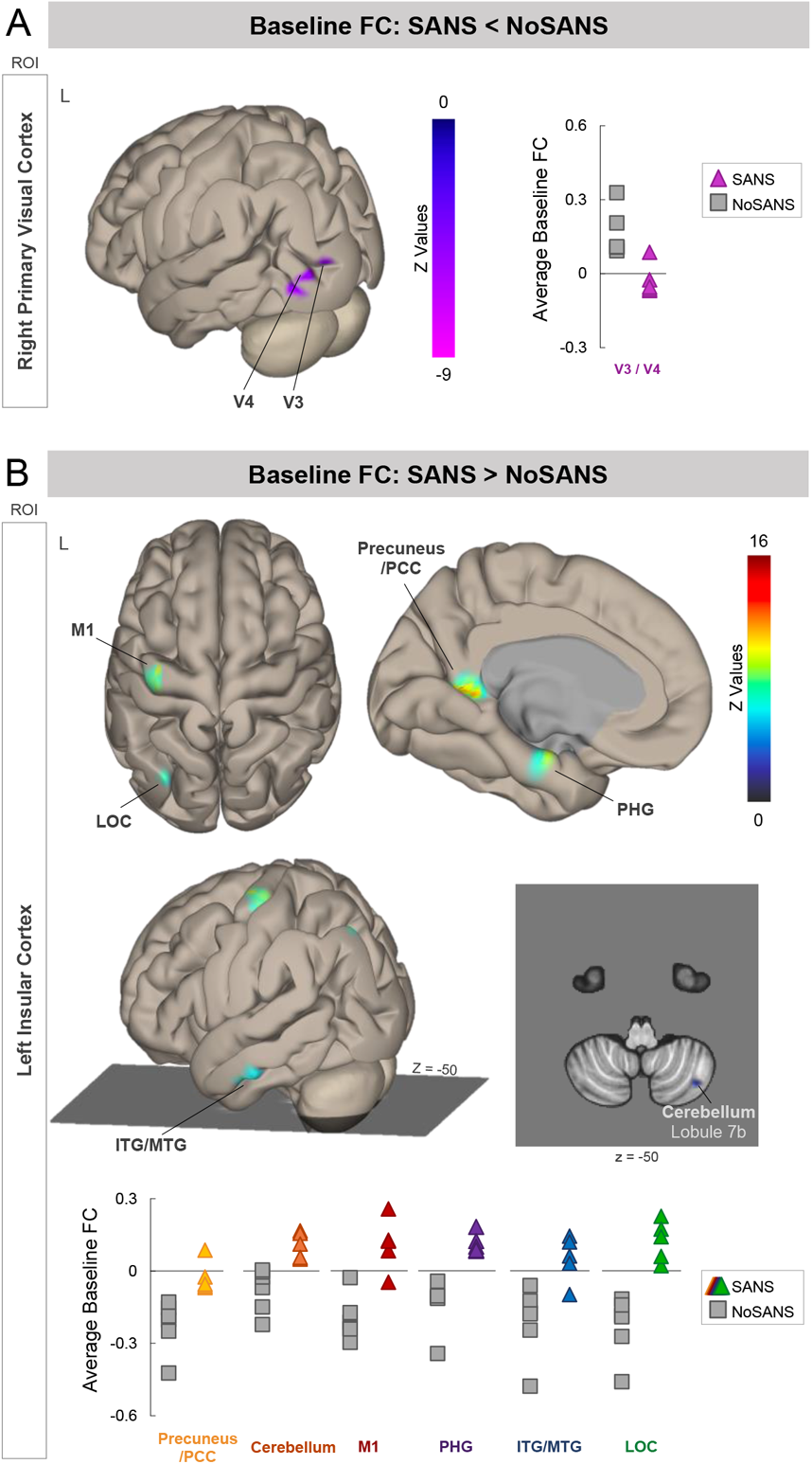
Pre-bed rest FC differences between the SANS and NoSANS subgroups. **A)** During the baseline phase, the NoSANS subgroup (gray squares) exhibited FC between the ROI in right primary visual cortex (V1) and visual areas V3 and V4 within the left hemisphere. The SANS subgroup exhibited an anti-correlation or lower FC between visual brain areas prior to bed rest (colored triangles) prior to the HDBR+CO_2_ intervention. **B)** During the baseline phase, the NoSANS subgroup (gray squares) exhibited anti-correlations between the ROI in left insular cortex and left primary motor cortex (M1), lateral occipital cortex (LOC), inferior and middle temporal gyri (ITG/MTG), parahippocampal gyrus (PHG), precuneus/posterior cingulate cortex (PCC), and right cerebellar lobule 7b. The SANS subgroup (colored triangles) exhibited greater FC than the NoSANS subgroup during the baseline phase. FC, functional connectivity; L, left; ROI, region of interest; SANS, spaceflight associated neuro-ocular syndrome.

**Table 5:** Peak MNI coordinates and T value of the cluster spanning visual areas V3 and V4 which exhibited lower functional connectivity with the ROI in primary visual cortex (V1) during the baseline phase for the SANS subgroup compared to the NoSANS subgroup. Cluster size indicates the number of voxels within each cluster. Uncorr cluster size p indicates uncorrected cluster-size p values. L, left; R, right; ROI, region of interest

| ROI | Target cluster | MNI coordinates |  |  | T value | Cluster size | Uncorr cluster size p |
| --- | --- | --- | --- | --- | --- | --- | --- |
|  |  | *x* | *y* | *z* |  |  |  |
| R primary visual cortex (V1) | L visual areas V3 / V4 | -38 | -88 | 2 | -11.83 | 125 | 0.000000009 |

The second network, identified using left insular cortex as a seed region, encompassed left primary motor cortex (M1), lateral occipital cortex (LOC), inferior and middle temporal gyri (ITG/MTG), parahippocampal gyrus (PHG), and right cerebellar lobule 7b. As shown in Figure 4B, the NoSANS subgroup exhibited FC within this network during the baseline phase. The SANS subgroup exhibited lower FC or an anti-correlation (i.e., negative FC) within this network during the baseline phase. Cluster coordinates and statistics are presented in Table 6.

**Table 6:** Peak MNI coordinates and T values of clusters which exhibited greater functional connectivity with the ROI in left insular cortex during the baseline phase for the SANS subgroup compared to the NoSANS subgroup. Cluster size indicates the number of voxels within each cluster. Uncorr cluster size p indicates uncorrected cluster-size p values. L, left; R, right; ROI, region of interest

| ROI | Target cluster | MNI coordinates |  |  | T value | Cluster size | Uncorr cluster size p |
| --- | --- | --- | --- | --- | --- | --- | --- |
|  |  | *x* | *y* | *z* |  |  |  |
| L insular cortex | L posterior cingulate (PCC) /precuneus | -4 | -46 | 2 | 15.73 | 125 | 0.00000009 |
|  | L primary motor cortex (M1) | -36 | -16 | 68 | 14.80 | 96 | 0.000001 |
|  | R cerebellum lobule 7b | 36 | -70 | -50 | 9.67 | 81 | 0.000004 |
|  | L parahippocampal gyrus (PHG) | -18 | -12 | -26 | 10.70 | 73 | 0.000009 |
|  | L lateral occipital cortex (LOC) | -32 | -80 | 50 | 9.54 | 53 | 0.000083 |
|  | L inferior temporal gyrus (ITG) | -54 | -18 | -24 | 7.21 | 47 | 0.000171 |

Our hypothesis-free voxel-to-voxel analysis did not yield any statistically significant clusters.

To assess the reliability of FC values between BDC-13 and BDC-7, we computed the intraclass correlation coefficient for each unique ROI-cluster connection for the networks shown in Figure 4. Table 7 shows the intraclass correlation coefficients and 95% confidence intervals for each unique ROI-cluster connection. For the network shown in Figure 4A, the intraclass correlation coefficient analysis yielded substantial test-retest reliability. For the network shown in Figure 4B, intraclass correlation coefficient analyses yielded fair (0.2-0.4) to almost perfect (>0.8) test-retest reliability across baseline sessions although confidence intervals were large. Figure 5 shows FC values from BDC-13 and BDC-7 sessions for each cluster within the networks shown in Figure 4.

**Table 7:** Average intraclass correlation coefficients for FC values between each ROI and target cluster and the 95% confidence interval (CI). ROI, region of interest; L, left; R, right

| ROI | Target cluster | Intraclass correlation coefficient | 95% CI |  |
| --- | --- | --- | --- | --- |
| L insular cortex | L posterior cingulate (PCC) /precuneus | 0.382 | -0.387 | 0.714 |
|  | L primary motor cortex (M1) | 0.833 | 0.351 | 0.956 |
|  | L parahippocampal gyrus (PHG) | 0.728 | -0.055 | 0.928 |
|  | R cerebellum lobule 7b | 0.687 | -0.253 | 0.917 |
|  | L lateral occipital cortex (LOC) | 0.817 | 0.292 | 0.951 |
|  | L inferior temporal gyrus (ITG) | 0.373 | -1.792 | 0.838 |
| R Primary visual cortex (V1) | L visual areas V3/V4 | 0.687 | -0.264 | 0.917 |

**Figure 5.**
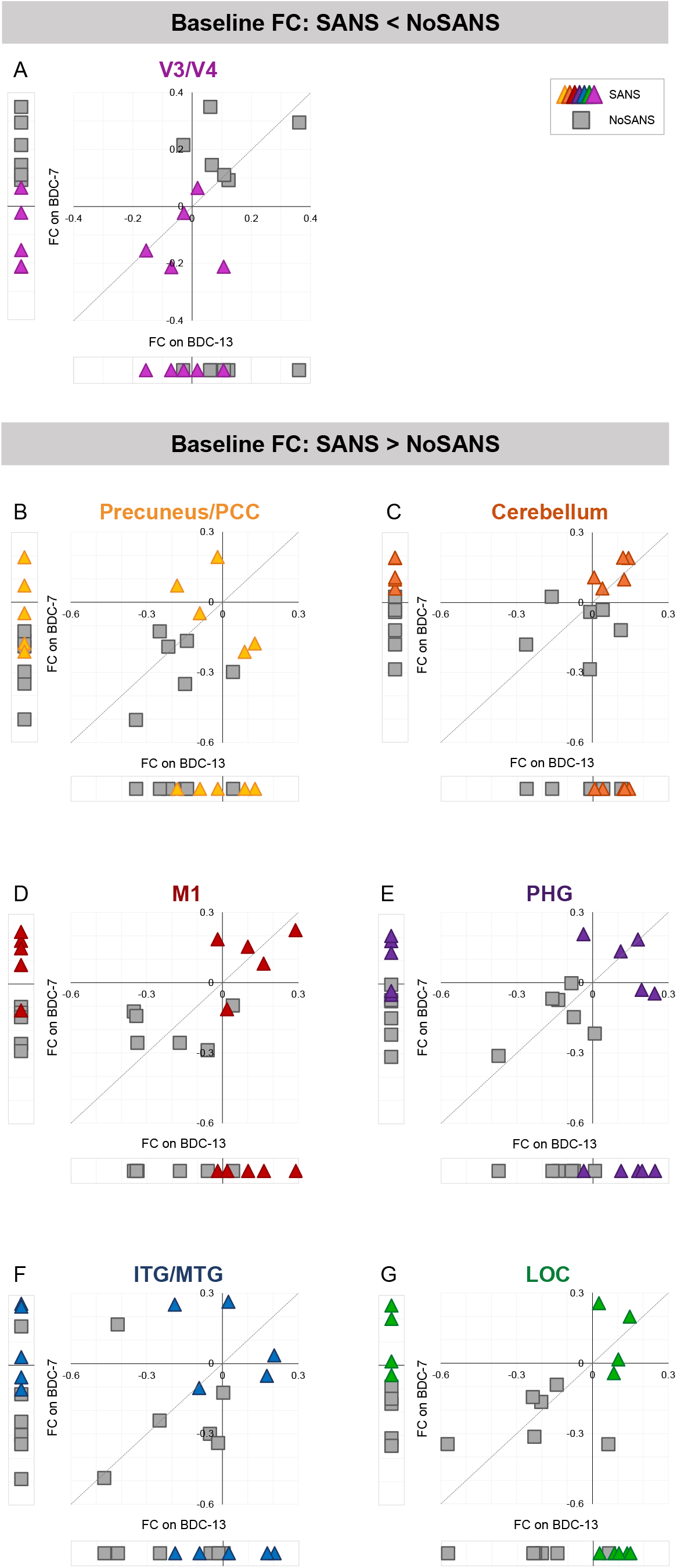
Baseline FC values. Within each panel, we present session-wise FC values between the indicated target cluster and the ROI in right primary visual cortex (A) or the ROI in left insular cortex (B-G). Each marker represents data from one participant. Triangle markers indicate participants who developed signs of SANS during the HDBR+CO_2_ intervention (SANS). Square markers indicated participants who did not develop signs of SANS (NoSANS). At the bottom of each scatterplot, we present subject-wise FC values from the BDC-13 session. To the left, we present subject-wise FC values estimated during the BDC-7 session. The scatterplot shows BDC-13 FC values plotted against BDC-7 FC values for each participant. The identity line is represented by the dashed line. FC, functional connectivity; BDC, baseline data collection; PCC, posterior cingulate cortex; M1, primary motor cortex; PHG, parahippocampal gyrus; ITG/MTG, inferior and middle temporal gyri; LOC, lateral occipital cortex; SANS, spaceflight associated neuro-ocular syndrome

### 3.3 Associations between Functional Connectivity and Genetic and B vitamin status

#### 3.3.1 Subgroup Differences in Longitudinal Changes in Functional Connectivity

In Figure 6, we show relationships between previously identified genetic and biochemical predictors of SANS and participants’ FC changes from pre- to post-HDBR+CO_2_ within the networks identified by our analyses of subgroup differences in longitudinal FC changes (Figure 3). Due to the known relationship between these biomarkers and SANS development within the current dataset ^18,19^, we have refrained from performing quantitative analyses. We present these associations for reference in future investigations.

**Figure 6.**
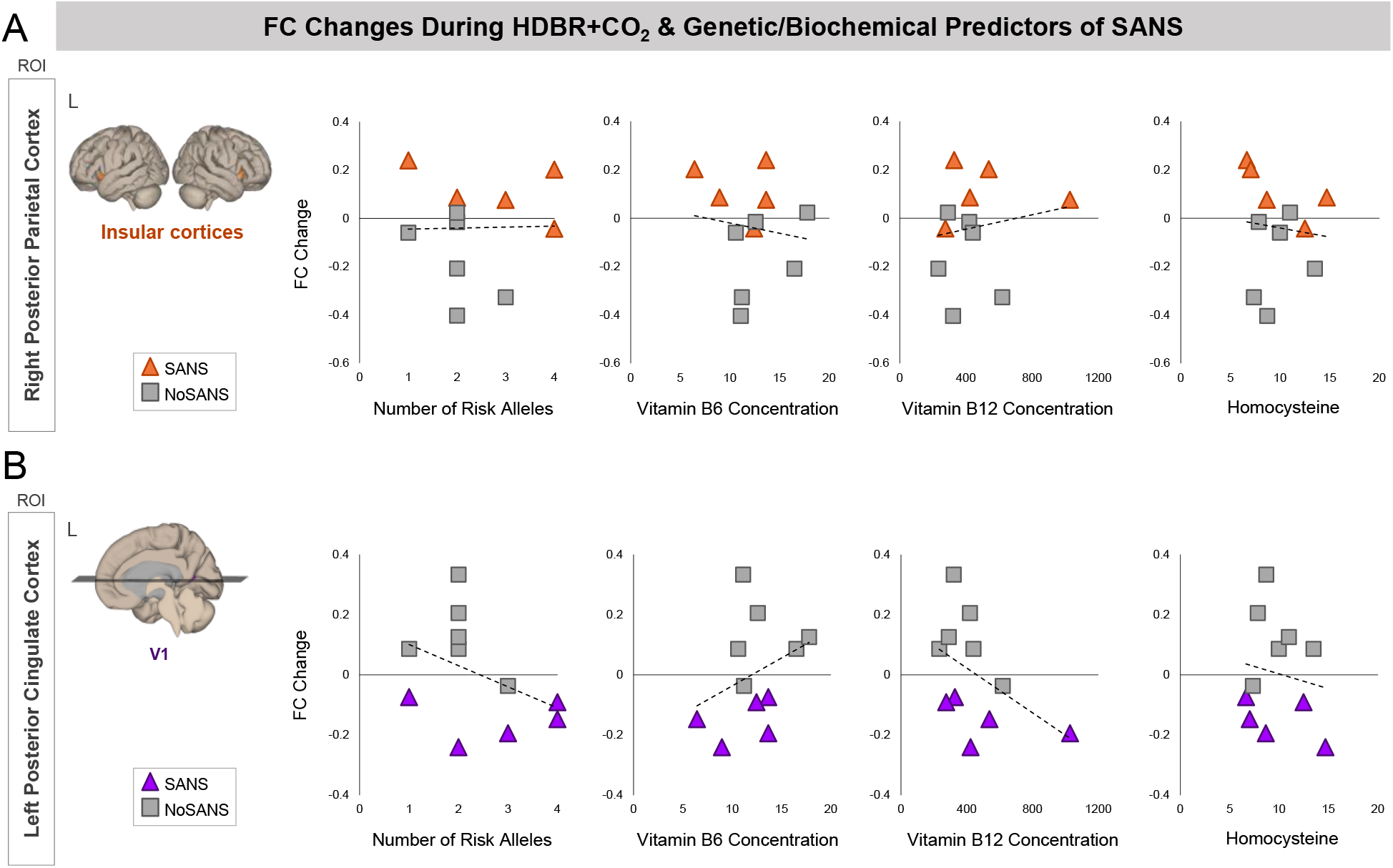
Associations between pre- to post-HDBR+CO_2_ FC values and SANS biomarkers. Scatterplots show relationships between the number of risk alleles, Vitamin B6 and B12 concentration, and homocysteine with participants’ change in FC from pre- to post-HDBR+CO_2_ (i.e., BDC-7 to HDT29). Triangle markers indicate participants who developed signs of SANS during the HDBR+CO_2_ intervention (SANS). Square markers indicated participants who did not develop signs of SANS (NoSANS). Dashed lines indicate the linear trendline across all 11 participants. **A)** In the upper panel, the y axes of plots indicate FC changes between the ROI in right posterior parietal cortex and target clusters in bilateral insular cortices. **B)** In the lower panel, the y axes of plots indicate FC changes between the ROI in left posterior cingulate cortex and target clusters in right primary visual cortex (V1). FC, functional connectivity; L, left; ROI, region of interest; SANS, spaceflight associated neuro-ocular syndrome.

#### 3.3.2 Subgroup Differences in Functional Connectivity at Baseline

In Figure 7, we show relationships between previously identified genetic and biochemical predictors of SANS and participants’ baseline FC values within the networks identified by our analyses of subgroup differences in baseline FC (Figure 4). The network identified using left insular cortex as an ROI consisted of six target clusters (Figure 4B). Here we have plotted associations between baseline FC and SANS biomarkers for one of the identified clusters. We selected the cluster spanning left precuneus/posterior cingulate cortex as its associations were representative of those of other clusters within the network.

**Figure 7.**
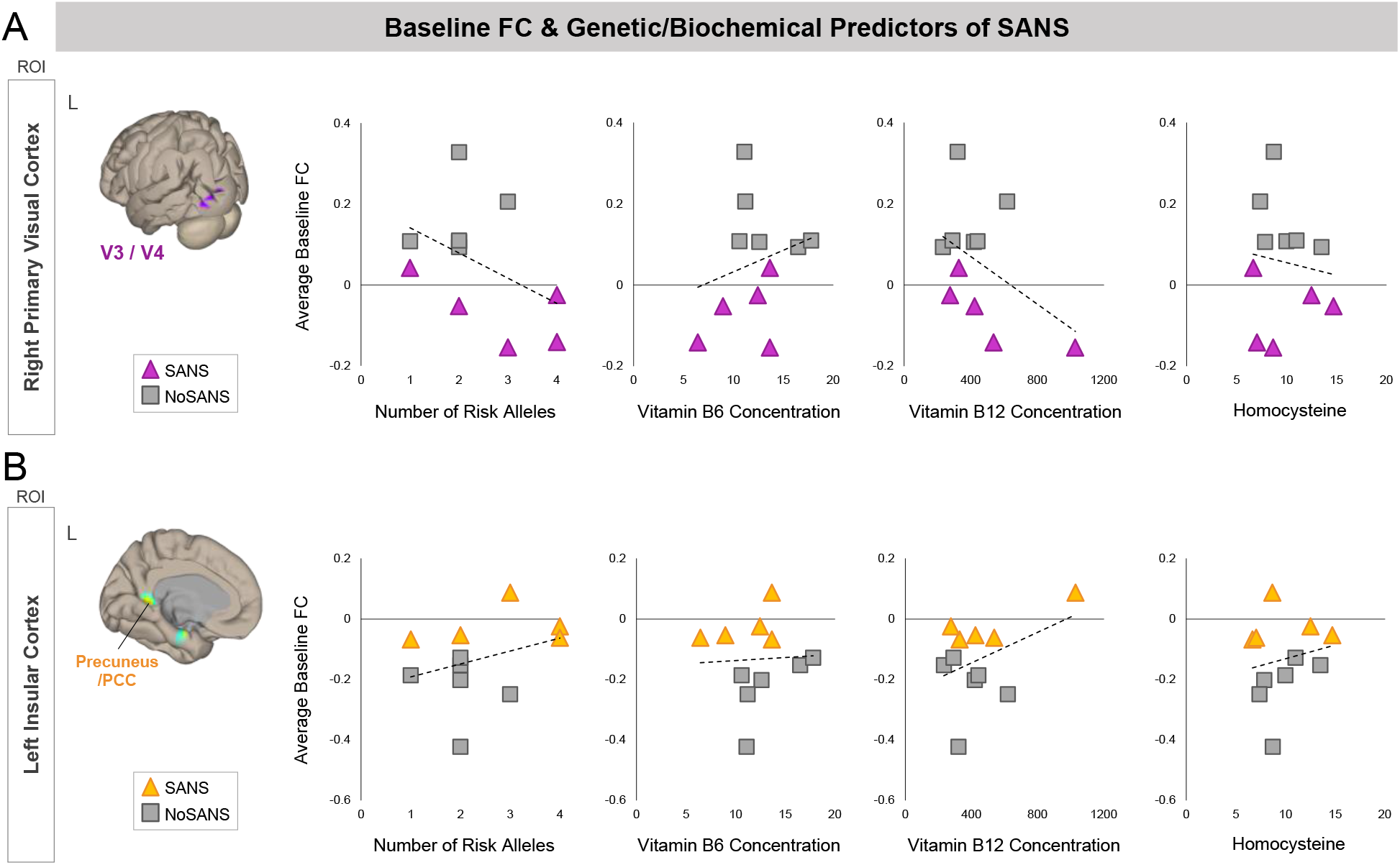
Associations between baseline FC values and SANS biomarkers. Scatterplots show relationships between the number of risk alleles, Vitamin B6 and B12 concentration, and homocysteine with participants’ pre-bed rest FC (i.e., averaged across BDC-13 and BDC-7). Triangle markers indicate participants who developed signs of SANS during the HDBR+CO_2_ intervention (SANS). Square markers indicated participants who did not develop signs of SANS (NoSANS). Dashed lines indicate the linear trendline across all 11 participants. **A)** In the upper panel, y axes indicate baseline FC values between the ROI in right primary visual cortex and target clusters spanning left V3 and V4. **B)** In the lower panel, y axes indicate baseline FC values between the ROI in left insular cortex and A target cluster in left precuneus/posterior cingulate cortex. FC, functional connectivity; L, left; ROI, region of interest; SANS, spaceflight associated neuro-ocular syndrome.

## 4. DISCUSSION

### 4.1 Subgroup Differences in Longitudinal Changes in Functional Connectivity

Here we show that the development of signs of SANS during a spaceflight analog is associated with a distinct pattern of resting-state FC changes involving visual and vestibular brain regions.

During the HDBR+CO_2_ intervention, the SANS subgroup exhibited an increase in FC between PPC and the insular cortices which reversed post-bed rest. The NoSANS subgroup exhibited the opposite pattern, showing FC decreases during HDBR+CO_2_ which reversed following bed rest. Posterior parietal cortex (PPC) is a region in which multisensory integration occurs, combining inputs from the visual, somatosensory and auditory systems ^49^. The insula is a region within the vestibular system ^50^, which uses multisensory integration for self-motion perception, spatial orientation, gaze stabilization, and postural control ^51^. During spaceflight and HDBR, the vestibular system receives altered or reduced sensory inputs and it is thought that the brain adaptively reweights sensory inputs as a result ^52–54^. The brain increases weighting of reliable sensory inputs or use in guiding performance while decreasing weighting of altered or unreliable inputs ^55^. The current results suggest differential patterns of vestibular reweighting during the HDBR+CO_2_ between the SANS and NoSANS subgroups. The SANS subgroup’s coupling between PPC and the insula may reflect progressively increased weighting of vestibular inputs during the spaceflight analog as optic disc edema develops. However, we did not find subgroup differences in vestibular processing in response to pneumatic skull taps from pre- to post-HDBR+CO_2_ ^56^. Future studies incorporating behavioral metrics of vestibular input reliance are required to investigate the functional significance of this finding.

The SANS subgroup exhibited reduced FC between posterior cingulate cortex (PCC) and primary visual cortex (V1) during HDBR+CO_2_ and then recovered post-bed rest whereas the NoSANS subgroup exhibited FC increases between these regions during HDBR+CO_2_. PCC is a key region within the default mode network. Although its function remains a topic of debate, recent theories suggest that the PCC is involved in task-independent introspection and self-referential processes ^57^. V1 is the first cortical stage of the visual system wherein low-level visual feature processing occurs ^58^. The differential patterns of FC change during HDBR+CO_2_ are likely a functional consequence of optic disc edema development. One hypothesis is that swelling of the optic disc exerts pressure on retinal ganglion cell axons as they exit the retina, subtly perturbing visual input signals en route to V1 (without impairing vision). The brain then responds by reducing the functional coupling between V1 and PCC, and by increasing the weighting on vestibular inputs. This hypothesis is speculative; future investigations are required to integrate visual and ophthalmic measures with assessments of V1 functional organization.

Notably, recent research has demonstrated spaceflight-related FC changes involving a subset of those regions identified by our longitudinal analyses, namely the insula and posterior cingulate cortex. A case study of a single cosmonaut showed decreases in intrinsic connectivity within the right insula and within PCC from pre- to post-flight ^59^. Moreover, a task-based connectivity study showed post-flight connectivity increases between the left and right insular cortices during plantar stimulation ^60^. These studies have also provided evidence for vestibular cortex reorganization and multisensory reweighting following spaceflight. The authors did not report if the cosmonauts who participated in these studies developed SANS during spaceflight, though it has been reported that cosmonauts do not develop SANS perhaps due to differences in in-flight countermeasures used by astronauts and cosmonauts^61^.

As CO_2_ is a potent arterial vasodilator, it is not possible to distinguish CO_2_-induced changes in vascular function from changes in neural activity on the basis of BOLD fMRI alone ^62^. Throughout this study, arterial partial pressure of carbon dioxide, assessed as part of NASA’s standard measures, showed a small but significant increase during the HDBR+CO_2_ phase ^16^ (though see also ^17,63^). Cerebral perfusion was also assessed during our MRI sessions using arterial spin labelling ^64^. As reported by Roberts et al ^64^, both subgroups showed a reduction in average perfusion across the whole brain during the HDBR+CO_2_ phase relative to baseline measures. On HDT7, the SANS subgroup exhibited a greater reduction in perfusion compared to the NoSANS subgroup. On HDT29, average cerebral perfusion of the SANS group exhibited partial recovery towards baseline levels whereas cerebral perfusion of the NoSANS subgroup remained consistent through the HDBR+CO_2_ phase. Perfusion values for both subgroups gradually returned towards baseline levels during the recovery phase, but remained 3% below baseline levels on R+12 ^64^. These cerebral perfusion results point to subgroup differences in vasoreactivity and autoregulatory responses during the HDBR+CO_2_ intervention. Subgroup differences in cerebral perfusion changes during the intervention may contribute to the longitudinal pattern FC changes presented here. However, drawing parallels between perfusion changes and regional FC changes during HDBR+CO_2_ is not clear as perfusion was assessed at the average across the entire brain ^64^. To address this issue, a follow-up investigation should assess cerebral perfusion changes within the regions constituting the FC networks identified in the current study.

As reported previously ^27^, the SANS and NoSANS subgroups exhibited differential patterns of performance change during the HDBTR+CO_2_ intervention. It’s not clear whether the FC changes during HDBR+CO_2_ would induce behavioural changes. Instead, these FC changes would be a marker of sensorimotor changes during the spaceflight analog. We think that the FC changes during bed rest are a marker of the brain’s sensory reweighting (i.e., relying more on one form of sensory input to make up for reductions/unreliability in another sensory modality). Sensory weighting and adaptation would likely be the driving forces behind performance changes during HDBR+CO_2_, and would be accompanied by FC changes.

### 4.2 Subgroup Differences in Functional Connectivity Before HDBR+CO_2_

We also identified two functional networks in which FC during the baseline phase was predictive of which participants would subsequently develop signs of SANS during HDBR+CO_2_. Participants who exhibited higher FC between the ROI in right primary visual cortex (V1) and left visual areas V3 and V4 during the baseline phase were less likely to develop signs of SANS during HDBR+CO_2_. Participants who lacked this coupling between primary and higher order visual areas during the baseline phase were more likely to develop signs of SANS during HDBR+CO_2_. V1 processes low-level visual stimuli ^58^ while higher level visual processing subserving visual recognition and visual attention are believed to occur within visual areas V3 and V4 ^65^.

We further identified a functional network consisting of left insula, primary motor cortex, lateral occipital cortex, inferior and middle temporal gyri, parahippocampal gyrus, precuneus/posterior cingulate cortex, and right cerebellar lobule 7b. Participants who exhibited greater decoupling between the insula and the other regions within the network during the baseline phase were less likely to develop signs of SANS during HDBR+CO_2_. In contrast, participants who exhibited coupling among these regions during the baseline phase were more likely to develop signs of SANS during HDBR+CO_2_. Many of the brain areas within this network support cognitive functions including visual perception, visuospatial processing, memory, and navigation ^66–68^.

These subgroup differences in pre-bed rest FC are likely markers of other differences that make subjects in the SANS subgroup more prone to developing optic disc edema. For example, recent work has identified genetic and biochemical predictors of optic disc edema development in the SANS subgroup of this study ^18,19^ as well as in astronauts who develop SANS ^20^. Low levels of B vitamins (B6, B12, riboflavin, and folate) and 1-carbon pathway risk alleles (all of which are linked to circulating homocysteine concentration) predict optic disc edema during HDBR+CO_2_ ^18,19^ and spaceflight ^21,69^. There is currently little evidence that individual differences in genetics and/or B vitamin status directly relate to functional connectivity^70^. Genetics, B vitamin status, and functional connectivity have all been linked to individual differences in anatomical connectivity. Lower vitamin B12 status and higher homocysteine concentration are associated with lower white matter volume in healthy older adults ^71,72^. Functional connectivity also shows a close correspondence to anatomical connectivity ^73^. It is thus feasible that subgroup differences in B vitamin levels and risk alleles in the current study ^18,19^ contribute to anatomical connectivity differences within the identified networks at baseline, which in turn result in individual differences in functional connectivity. However, if this were the case, one would expect the SANS subgroup to exhibit lower FC compared to the NoSANS group for both networks. Inclusion of anatomical brain connectivity measures are required to directly test this hypothesis.

Analyses of average whole brain cerebral perfusion in these same subjects have shown no significant differences between the SANS and NoSANS subgroups prior to bed rest ^64^. However, since perfusion analyses were performed at the whole-brain level, they may not have captured regional differences in perfusion between the subgroups.

Future research is required to further investigate the cause(s) of SANS, and whether or how these pre-bed rest FC differences relate to the cause(s). Direct regional comparisons of structural, perfusion, and resting-state fMRI data between the SANS and NoSANS groups during the baseline phase are required to disentangle the origin of the subgroup FC differences within the FC networks reported here.

### 4.3 Limitations

The small sample size is a significant limitation of our work. This study was the first long-duration bed rest study to implement strict HDBR and use a mild hypercapnic environment. As such, we are unable to determine whether the development of optic disc edema was due to strict HDBR, elevated CO_2_ levels, or a combination thereof. Moreover, optic disc edema in the SANS subgroup was more pronounced than typically seen in astronauts post-flight ^3^. In addition, cases of SANS in astronauts typically include other ophthalmic changes including globe flattening, choroidal folds, cotton wool spots, etc. ^1^. Thus, it remains unclear how closely the signs of SANS observed in the current dataset mimic those seen in astronauts.

Visual deterioration during long-duration spaceflight represents a significant potential risk to human health and performance on current and future missions. Therefore, despite the limitations noted above, these data have provided important insights into the underlying cause of SANS ^21,69^ and understanding the breadth of its effects. This is particularly the case given that prior work has only related SANS to central nervous system structures, and not to connectivity or activity.

### 4.4 Conclusion

Here we have shown that the SANS and NoSANS subgroups exhibited differential resting-state brain activity prior to the spaceflight analog within a visual cortical network and within a large-scale network of brain areas involved in multisensory integration. We also showed that the development of signs of SANS during the spaceflight analog is associated with differential patterns of resting-state connectivity changes within visual and vestibular-related brain networks. This finding suggests that SANS impacts not only neuro-ocular structures, but also brain function. Forthcoming investigations by our group will combine functional and structural brain analyses to assess whether astronauts who develop SANS exhibit similar patterns of pre-flight FC and/or pre- to post-flight FC changes.

## REFERENCES

1. Mader TH, Gibson CR, Pass AF, et al. Optic disc edema, globe flattening, choroidal folds, and hyperopic shifts observed in astronauts after long-duration space flight. Ophthalmology. 2011;118(10):2058–2069.

2. Lee AG, Mader TH, Gibson CR, et al. Spaceflight associated neuro-ocular syndrome (SANS) and the neuro-ophthalmologic effects of microgravity: a review and an update. NPJ Microgravity. 2020;6:7.

3. Laurie SS, Lee SMC, Macias BR, et al. Optic Disc Edema and Choroidal Engorgement in Astronauts During Spaceflight and Individuals Exposed to Bed Rest. JAMA Ophthalmol. Published online December 26, 2019. doi:10.1001/jamaophthalmol.2019.5261

4. Pavy-Le Traon A, Heer M, Narici MV, Rittweger J, Vernikos J. From space to Earth: advances in human physiology from 20 years of bed rest studies (1986-2006). Eur J Appl Physiol. 2007;101(2):143–194.

5. Cassady K, Koppelmans V, Reuter-Lorenz P, et al. Effects of a spaceflight analog environment on brain connectivity and behavior. Neuroimage. 2016;141:18–30.

6. Kramer LA, Hasan KM, Stenger MB, et al. Intracranial Effects of Microgravity: A Prospective Longitudinal MRI Study. Radiology. 2020;295(3):640–648.

7. Alperin N, Bagci AM. Spaceflight-Induced Visual Impairment and Globe Deformations in Astronauts Are Linked to Orbital Cerebrospinal Fluid Volume Increase. Acta Neurochir Suppl. 2018;126:215–219.

8. Van Ombergen A, Jillings S, Jeurissen B, et al. Brain ventricular volume changes induced by long-duration spaceflight. Proc Natl Acad Sci U S A. 2019;116(21):10531–10536.

9. Taibbi G, Cromwell RL, Zanello SB, et al. Ocular Outcomes Comparison Between 14-and 70-Day Head-Down-Tilt Bed Rest. Invest Ophthalmol Vis Sci. 2016;57(2):495–501.

10. Stenger MB, Tarver WJ, Brunstetter T, Gibson CR, Laurie SS and Lee, Stuart and Macias, Brandon R and Mader, Thomas H and Otto Christian and Smith, Scott M. Evidence report: risk of Spaceflight Associated Neuro-ocular Syndrome (SANS). Published online 2017.

11. Law J, Van Baalen M, Foy M, et al. Relationship between carbon dioxide levels and reported headaches on the international space station. J Occup Environ Med. 2014;56(5):477–483.

12. Artru AA. Reduction of cerebrospinal fluid pressure by hypocapnia: changes in cerebral blood volume, cerebrospinal fluid volume, and brain tissue water and electrolytes. J Cereb Blood Flow Metab. 1987;7(4):471–479.

13. Moseley ME, Chew WM, White DL, et al. Hypercarbia-induced changes in cerebral blood volume in the cat: A1H MRI and intravascular contrast agent study. Magnetic Resonance in Medicine. 1992;23(1):21–30. doi:10.1002/mrm.1910230104

14. Fortune JB, Feustel PJ, deLuna C, Graca L, Hasselbarth J, Kupinski AM. Cerebral blood flow and blood volume in response to O2 and CO2 changes in normal humans. J Trauma. 1995;39(3):463-471; discussion 471-472.

15. Lawley JS, Petersen LG, Howden EJ, et al. Effect of gravity and microgravity on intracranial pressure. J Physiol. 2017;595(6):2115–2127.

16. McGregor HR, Lee JK, Mulder ER, et al. Brain connectivity and behavioral changes in a spaceflight analog environment with elevated CO2. NeuroImage. 2020;225:117450. doi:10.1016/j.neuroimage.2020.117450

17. Laurie SS, Macias BR, Dunn JT, et al. Optic Disc Edema after 30 Days of Strict Head-down Tilt Bed Rest. Ophthalmology. 2019;126(3):467–468.

18. Zwart SR, Laurie SS, Chen JJ, et al. Association of Genetics and B Vitamin Status With the Magnitude of Optic Disc Edema During 30-Day Strict Head-Down Tilt Bed Rest. JAMA Ophthalmology. 2019;137(10):1195. doi:10.1001/jamaophthalmol.2019.3124

19. Smith S, Laurie S, Young M, Zwart S. MTRR 66 and SHMT1 1420 Variants Are Associated with Optic Disc Edema During 30-d Strict Head-down Tilt Bed Rest and CO2 Exposure (P24-036-19). Current Developments in Nutrition. 2019;3(Supplement_1). doi:10.1093/cdn/nzz044.p24-036-19

20. Zwart SR, Gregory JF, Zeisel SH, et al. Genotype, B-vitamin status, and androgens affect spaceflight-induced ophthalmic changes. FASEB J. 2016;30(1):141–148.

21. Smith SM, Zwart SR. Spaceflight-related ocular changes: the potential role of genetics, and the potential of B vitamins as a countermeasure. Curr Opin Clin Nutr Metab Care. 2018;21(6):481–488.

22. Koppelmans V, Bloomberg JJ, De Dios YE, et al. Brain plasticity and sensorimotor deterioration as a function of 70 days head down tilt bed rest. PLoS One. 2017;12(8):e0182236.

23. Liao Y, Zhang J, Huang Z, et al. Altered Baseline Brain Activity with 72 h of Simulated Microgravity – Initial Evidence from Resting-State fMRI. PLoS ONE. 2012;7(12):e52558. doi:10.1371/journal.pone.0052558

24. Liao Y, Lei M, Huang H, et al. The time course of altered brain activity during 7-day simulated microgravity. Front Behav Neurosci. 2015;9:124.

25. Zhou Y, Wang Y, Rao L-L, et al. Disrupted resting-state functional architecture of the brain after 45-day simulated microgravity. Frontiers in Behavioral Neuroscience. 2014;8. doi:10.3389/fnbeh.2014.00200

26. Demertzi A, Van Ombergen A, Tomilovskaya E, et al. Cortical reorganization in an astronaut’s brain after long-duration spaceflight. Brain Struct Funct. 2016;221(5):2873–2876.

27. Lee JK, De Dios Y, Kofman I, Mulavara AP, Bloomberg JJ, Seidler RD. Head Down Tilt Bed Rest Plus Elevated CO2 as a Spaceflight Analog: Effects on Cognitive and Sensorimotor Performance. Frontiers in Human Neuroscience. 2019;13. doi:10.3389/fnhum.2019.00355

28. Laurie SS, Christian K, Kysar J, et al. Unchanged cerebrovascular CO reactivity and hypercapnic ventilatory response during strict head-down tilt bed rest in a mild hypercapnic environment. J Physiol. 2020;598(12):2491–2505.

29. Whitfield-Gabrieli S, Nieto-Castanon A. Conn: a functional connectivity toolbox for correlated and anticorrelated brain networks. Brain Connect. 2012;2(3):125–141.

30. Romero JE, Coupé P, Giraud R, et al. CERES: A new cerebellum lobule segmentation method. NeuroImage. 2017;147:916–924. doi:10.1016/j.neuroimage.2016.11.003

31. Carass A, Cuzzocreo JL, Han S, et al. Comparing fully automated state-of-the-art cerebellum parcellation from magnetic resonance images. Neuroimage. 2018;183:150–172.

32. Diedrichsen J. A spatially unbiased atlas template of the human cerebellum. Neuroimage. 2006;33(1):127–138.

33. Van Dijk KRA, Sabuncu MR, Buckner RL. The influence of head motion on intrinsic functional connectivity MRI. NeuroImage. 2012;59(1):431–438. doi:10.1016/j.neuroimage.2011.07.044

34. Power JD, Barnes KA, Snyder AZ, Schlaggar BL, Petersen SE. Spurious but systematic correlations in functional connectivity MRI networks arise from subject motion. Neuroimage. 2012;59(3):2142–2154.

35. Van Dijk Kra, Hedden T, Venkataraman A, Evans KC, Lazar SW, Buckner RL. Intrinsic functional connectivity as a tool for human connectomics: theory, properties, and optimization. J Neurophysiol. 2010;103(1):297–321.

36. Cole DM, Smith SM, Beckmann CF. Advances and pitfalls in the analysis and interpretation of resting-state FMRI data. Front Syst Neurosci. 2010;4:8.

37. Behzadi Y, Restom K, Liau J, Liu TT. A component based noise correction method (CompCor) for BOLD and perfusion based fMRI. Neuroimage. 2007;37(1):90–101.

38. Biswal B, Yetkin FZ, Haughton VM, Hyde JS. Functional connectivity in the motor cortex of resting human brain using echo-planar MRI. Magn Reson Med. 1995;34(4):537–541.

39. Damoiseaux JS, S A R Barkhof F, et al. Consistent resting-state networks across healthy subjects. Proceedings of the National Academy of Sciences. 2006;103(37):13848–13853.

40. Zeng L-L, Liao Y, Shen H, Liu X, Hu D. Decoding Brain States with Simulated Microgravity from Baseline Using Functional Connectivity of Default Network. Advances in Cognitive Neurodynamics (V). Published online 2016:325-330. doi:10.1007/978-981-10-0207-6_45

41. Zeng L-L, Liao Y, Zhou Z, et al. Default network connectivity decodes brain states with simulated microgravity. Cognitive Neurodynamics. 2016;10(2):113–120. doi:10.1007/s11571-015-9359-8

42. Liao Y, Miao D, Huan Y, Yin H, Xi Y, Liu X. Altered regional homogeneity with short-term simulated microgravity and its relationship with changed performance in mental transformation. PLoS One. 2014;8(6):e64931.

43. Demertzi A, Van Ombergen A, Tomilovskaya E, et al. Cortical reorganization in an astronaut’s brain after long-duration spaceflight. Brain Struct Funct. 2016;221(5):2873–2876.

44. Martuzzi R, Ramani R, Qiu M, Shen X, Papademetris X, Todd Constable R. A whole-brain voxel based measure of intrinsic connectivity contrast reveals local changes in tissue connectivity with anesthetic without a priori assumptions on thresholds or regions of interest. NeuroImage. 2011;58(4):1044–1050. doi:10.1016/j.neuroimage.2011.06.075

45. Landis JR, Richard Landis J, Koch GG. The Measurement of Observer Agreement for Categorical Data. Biometrics. 1977;33(1):159. doi:10.2307/2529310

46. Benjamini Y, Hochberg Y. Controlling the False Discovery Rate: A Practical and Powerful Approach to Multiple Testing. Journal of the Royal Statistical Society: Series B (Methodological). 1995;57(1):289–300. doi:10.1111/j.2517-6161.1995.tb02031.x

47. Chumbley J, Worsley K, Flandin G, Friston K. Topological FDR for neuroimaging. Neuroimage. 2010;49(4):3057–3064.

48. Poldrack RA, Mumford JA. Independence in ROI analysis: where is the voodoo? Social Cognitive and Affective Neuroscience. 2009;4(2):208–213. doi:10.1093/scan/nsp011

49. Lewis JW, Van Essen DC. Corticocortical connections of visual, sensorimotor, and multimodal processing areas in the parietal lobe of the macaque monkey. J Comp Neurol. 2000;428(1):112–137.

50. zu Eulenburg P, Caspers S, Roski C, Eickhoff SB. Meta-analytical definition and functional connectivity of the human vestibular cortex. Neuroimage. 2012;60(1):162–169.

51. Balaban CD. Neurotransmitters in the vestibular system. Handbook of Clinical Neurology. Published online 2016:41–55. doi:10.1016/b978-0-444-63437-5.00003-0

52. Young L, Oman C, Watt D, Money K, Lichtenberg B. Spatial orientation in weightlessness and readaptation to earth’s gravity. Science. 1984;225(4658):205–208. doi:10.1126/science.6610215

53. Young LR, Oman CM, Watt DG, et al. M.I.T./Canadian vestibular experiments on the Spacelab-1 mission: 1. Sensory adaptation to weightlessness and readaptation to one-g: an overview. Exp Brain Res. 1986;64(2):291–298.

54. Mulavara AP, Peters BT, Miller CA, et al. Physiological and Functional Alterations after Spaceflight and Bed Rest. Med Sci Sports Exerc. 2018;50(9):1961–1980.

55. Assländer L, Peterka RJ. Sensory reweighting dynamics in human postural control. J Neurophysiol. 2014;111(9):1852–1864.

56. Hupfeld KE, Lee JK, Gadd NE, et al. Neural Correlates of Vestibular Processing During a Spaceflight Analog With Elevated Carbon Dioxide (CO2): A Pilot Study. Frontiers in Systems Neuroscience. 2020;13. doi:10.3389/fnsys.2019.00080

57. Leech R, Sharp DJ. The role of the posterior cingulate cortex in cognition and disease. Brain. 2014;137(1):12–32. doi:10.1093/brain/awt162

58. Tootell RBH, Hadjikhani NK, Vanduffel W, et al. Functional analysis of primary visual cortex (V1) in humans. Proceedings of the National Academy of Sciences. 1998;95(3):811–817. doi:10.1073/pnas.95.3.811

59. Demertzi A, Van Ombergen A, Tomilovskaya E, et al. Cortical reorganization in an astronaut’s brain after long-duration spaceflight. Brain Struct Funct. 2016;221(5):2873–2876.

60. Pechenkova E, Nosikova I, Rumshiskaya A, et al. Alterations of Functional Brain Connectivity After Long-Duration Spaceflight as Revealed by fMRI. Front Physiol. 2019;10:761.

61. Jillings S, Van Ombergen A, Tomilovskaya E, et al. Macro-and microstructural changes in cosmonauts’ brains after long-duration spaceflight. Science Advances. 2020;6(36):eaaz9488. doi:10.1126/sciadv.aaz9488

62. Drew PJ. Vascular and neural basis of the BOLD signal. Curr Opin Neurobiol. 2019;58:61–69.

63. Laurie SS, Christian K, Kysar J, et al. Unchanged cerebrovascular CO 2 reactivity and hypercapnic ventilatory response during strict head-down tilt bed rest in a mild hypercapnic environment. The Journal of Physiology. 2020;598(12):2491–2505. doi:10.1113/jp279383

64. Roberts DR, Collins HR, Lee JK, et al. Altered cerebral perfusion in response to chronic mild hypercapnia and head-down tilt Bed rest as an analog for Spaceflight. Neuroradiology. Published online February 15, 2021. doi:10.1007/s00234-021-02660-8

65. Roe AW, Chelazzi L, Connor CE, et al. Toward a Unified Theory of Visual Area V4. Neuron. 2012;74(1):12–29. doi:10.1016/j.neuron.2012.03.011

66. Aminoff EM, Kveraga K, Bar M. The role of the parahippocampal cortex in cognition. Trends in Cognitive Sciences. 2013;17(8):379–390. doi:10.1016/j.tics.2013.06.009

67. Stoodley CJ, Schmahmann JD. Evidence for topographic organization in the cerebellum of motor control versus cognitive and affective processing. Cortex. 2010;46(7):831–844.

68. Aguirre GK, Singh R, D’Esposito M. Stimulus inversion and the responses of face and object-sensitive cortical areas. Neuroreport. 1999;10(1):189–194.

69. Zwart SR, Gibson CR, Gregory JF, et al. Astronaut ophthalmic syndrome. The FASEB Journal. 2017;31(9):3746–3756. doi:10.1096/fj.201700294

70. Downey LA, Simpson TN, Ford TC, et al. Increased Posterior Cingulate Functional Connectivity Following 6-Month High-Dose B-Vitamin Multivitamin Supplementation: A Randomized, Double-Blind, Placebo-Controlled Trial. Front Nutr. 2019;6:156.

71. Vogiatzoglou A, Refsum H, Johnston C, et al. Vitamin B12 status and rate of brain volume loss in community-dwelling elderly. Neurology. 2008;71(11):826–832.

72. Feng L, Isaac V, Sim S, Ng T-P, Krishnan KRR, Chee MWL. Associations between elevated homocysteine, cognitive impairment, and reduced white matter volume in healthy old adults. Am J Geriatr Psychiatry. 2013;21(2):164–172.

73. Fox MD, Snyder AZ, Vincent JL, Corbetta M, Van Essen DC, Raichle ME. The human brain is intrinsically organized into dynamic, anticorrelated functional networks. Proc Natl Acad Sci U S A. 2005;102(27):9673–9678.

